# Differential importance of MSP4 and MSP5 for infection of red blood cells between human infecting malaria parasites

**DOI:** 10.1101/2025.08.28.672823

**Authors:** Jill Chmielewski, Isabelle G. Henshall, Ornella Romeo, Danny W Wilson

## Abstract

*Plasmodium* species malaria parasites require invasion and replication within red blood cells to cause disease. Merozoite surface proteins (MSPs) are proposed to play a role in attachment of merozoites to RBCs and have long been considered as potential vaccine targets, but their functions during invasion are largely unknown. We applied targeted gene editing to investigate MSP4 and 5 function in *P. falciparum*, which causes most malaria mortality, and *P. knowlesi*, an *in vitro* culturable zoonotic species closely related to the widespread *P. vivax*. CRISPR-Cas9 gene-editing revealed that *P. knowlesi* MSP4 was not required for parasite growth *in vitro*. While *P. knowlesi* MSP5 could be functionally replaced by *P. vivax* MSP5, it was refractory to gene deletion. We confirmed the opposite for two different *P. falciparum* laboratory isolates where MSP4 is essential but MSP5 is dispensable. Attempts to select for reliance on the non-essential MSP (e.g. *P. knowlesi* MSP4 or *P. falciparum* MSP5) through long-term growth of inducible knock-out parasites, or via chimeric complementation of the essential MSP4 or 5 with the essential MSP from the other species, were unsuccessful. Live cell filming revealed a severe cell-entry defect with conditional knock-down of MSP5 protein expression in *P. knowlesi*. This study demonstrates differential importance of MSP4 and MSP5 during merozoite RBC invasion across human infecting malaria species, emphasises that vaccine candidates must be considered individually for the two most prominent human malarias and promotes MSP5 as a potential vaccine candidate for *P. knowlesi* and *P. vivax*.

**Significance:** For a malaria parasite to cause disease, the merozoite form of the lifecycle has to infect and replicate within human red blood cells. Proteins on the surface of the merozoite are considered as promising vaccine candidates, but the functions of these proteins are poorly understood. Here we demonstrate that two structurally similar merozoite surface proteins (MSP), MSP4 and MSP5, have differential importance between one human infecting malaria species compared to a second. The finding that MSP4 is essential for growth in one species, and MSP5 in the other, has implications for understanding invasion biology of malaria parasites and highlights that even structurally similar vaccine targets may need to be chosen specifically for each human infecting malaria species.

## Introduction

Malaria is a parasitic disease caused by *Plasmodium* species parasites. In 2023, the WHO estimated there were 263 million cases resulting in 597,000 deaths. This reflects a concerning trend in case and fatality numbers plateauing and even increasing in some countries (1, 2). Although we know that *P. falciparum* causes the majority of malaria mortality, there is significant morbidity caused by *P. vivax* outside of the African continent (3) and the inability to culture this species *in vitro* long term has made it difficult to study it’s biology. *P. knowlesi* is a zoonotic species which is an increasingly common cause of severe malaria (4, 5) and is more closely related to *P. vivax* than *P. falciparum. P. knowlesi* has been adapted to *in vitro* culture (6) and is amenable to gene editing (7, 8) and functional expression of *P. vivax* antigens, making it a useful laboratory tool for vaccine candidate screening (9).

The clinical symptoms of malaria are caused by the asexual blood stage replication cycle, which begins when a merozoite invades a red blood cell (RBC). After a period of growth, the parasite makes ∼10 – 30 RBC invading merozoites (depending on the species) which rupture the RBC and enter circulation where they then proceed to invade the next RBC, typically within minutes (10, 11). The biology of merozoite RBC entry is not entirely understood. However, it is known that proteins unique to malaria, and sometimes related parasites, which reside on the surface or are secreted from specialised organelles at the apical tip of the invasive merozoite have important roles in invasion.

Merozoite surface proteins (MSPs), named for their shared surface localisation and not necessarily because of any structural relationship, are either GPI anchored or peripherally associate in complexes and have been shown to be a major component of the protein content on the parasite surface (12). Their surface localisation places them as potentially the first point of contact between the parasite and the host RBC, leading to the assumption they include mediators of RBC invasion. MSPs have been identified as targets of naturally acquired immunity and some are targets of antibodies that are capable of inhibiting parasite replication *in vitro* (13–19). However, *in vitro* growth inhibition has also proven to be an unreliable predictor of clinical protection (20, 21).

Despite studies giving evidence to MSPs being targets of growth inhibitory antibodies, there is a dearth of direct functional evidence showing they have a role in RBC invasion. Initial gene-editing studies of MSP2 essentiality for *P. falciparum* reported *Pf*MSP1, *Pf*MSP2, *Pf*MSP4 and *Pf*MSP10 as essential for parasite growth *in vitro* (22). On the other hand, *Pf*MSP5 (22) and members of the *Pf*MSP3 and *Pf*MSP7 families (23, 24) could be knocked out with no substantial impact on *in vitro* growth.

With the adaptation of CRISPR-Cas9 gene editing for use in *Plasmodium* species (7, 25) the assumed role of MSPs in merozoite invasion is being re-examined and we are gaining new insights into some of the previously most well studied MSPs. Specifically, recent studies have revealed that the two most abundant MSPs of *P. falciparum,* MSP1 and MSP2, may have no direct role in merozoite RBC invasion. *Pf*MSP1, a major target of antibodies in malaria exposed samples, makes up ∼30% of the GPI anchored protein content on the surface of *P. falciparum* merozoites and was found to be refractory to gene deletion (12, 21, 22, 26, 27). Initially believed to have a role in merozoites interacting with the RBC surface (28), recent studies using an inducible *Pf*MSP1 knock-out line have demonstrated that *Pf*MSP1 is necessary for rupture of the schizont rather than merozoite invasion (29) and inducible knock-out of *Pf*MSP1 has no influence on the strength of binding to the RBC (30). A recent study by Henshall et al. (31) demonstrated *Pf*MSP2, the second most abundant MSP on the *P. falciparum* merozoite surface (12), can be knocked out with no change to *in vitro* parasite replication. As such, even though *Pf*MSP1 and *Pf*MSP2 are abundant proteins on the surface of the *P. falciparum* merozoite, the latest evidence suggests their role in RBC invasion may be limited.

Currently, there is little direct evidence that any other MSPs have a functional role in merozoite invasion of the RBC. The MSP1 bound MSP7 gene family is made up of paralogous genes which have duplicated to varying degrees in different *Plasmodium* lineages (32). A study by Kadekoppala et al. showed genetic deletion of the *P. falciparum* paralog *Pf*MSP7-1 (3D7_1335100) resulted in a modest ∼20% reduction in blood stage growth following a single replication cycle (23), but whether this was due to loss of merozoite invasion, or another growth defect, was not investigated. Importantly, gene-editing studies to investigate the function of MSPs in human malaria parasites are largely limited to *P. falciparum*. A single study for *P. knowlesi* in which the paralog of *Pk*MSP1, *Pk*MSP1P, was knocked out, showed that this protein is important for *in vitro* blood stage replication (33). These studies highlight that our understanding of MSP function is often crude and limited to one species of human malaria, and that application of newly developed and more efficient gene-editing techniques are needed to provide greater insights into MSP function.

MSP4 and MSP5 likely arose from a gene-duplication event and exist in a highly syntenic region of the genome of ape, monkey, and bird malaria parasites, with these genes represented by a single MSP4/5 fusion in rodent malaria parasites (34–37) (Figure 1, a, b, c). MSP4 and MSP5 are GPI anchored proteins with single EGF-like domains (34–37) and together account for ∼6% of the GPI-anchored protein content on the surface of the *P. falciparum* merozoite (12). Early investigation into function and immunogenicity of *Pf*MSP4 indicated it was antigenic (38) (Wang et al 1999) and refractory to genetic deletion, but *Pf*MSP5 was amenable to deletion with no effect on *in vitro* blood stage replication (22). Immunisation studies in mice using the dual MSP4/5 protein of *P. yoelii* reported vaccination with this antigen could induce protection from subsequent lethal parasite challenge (39). Studies of naturally occurring immunity in humans have shown an association between anti-*Pf*MSP4 antibodies and protection from future clinical episodes of malaria (18) and that both *P. falciparum* and *P. vivax* infection elicit anti-MSP5 antibodies (40). These studies support that MSP4 and MSP5 could play a role in providing protective immunity to either *P. falciparum* or *P. vivax* malaria.

**Figure 1.**
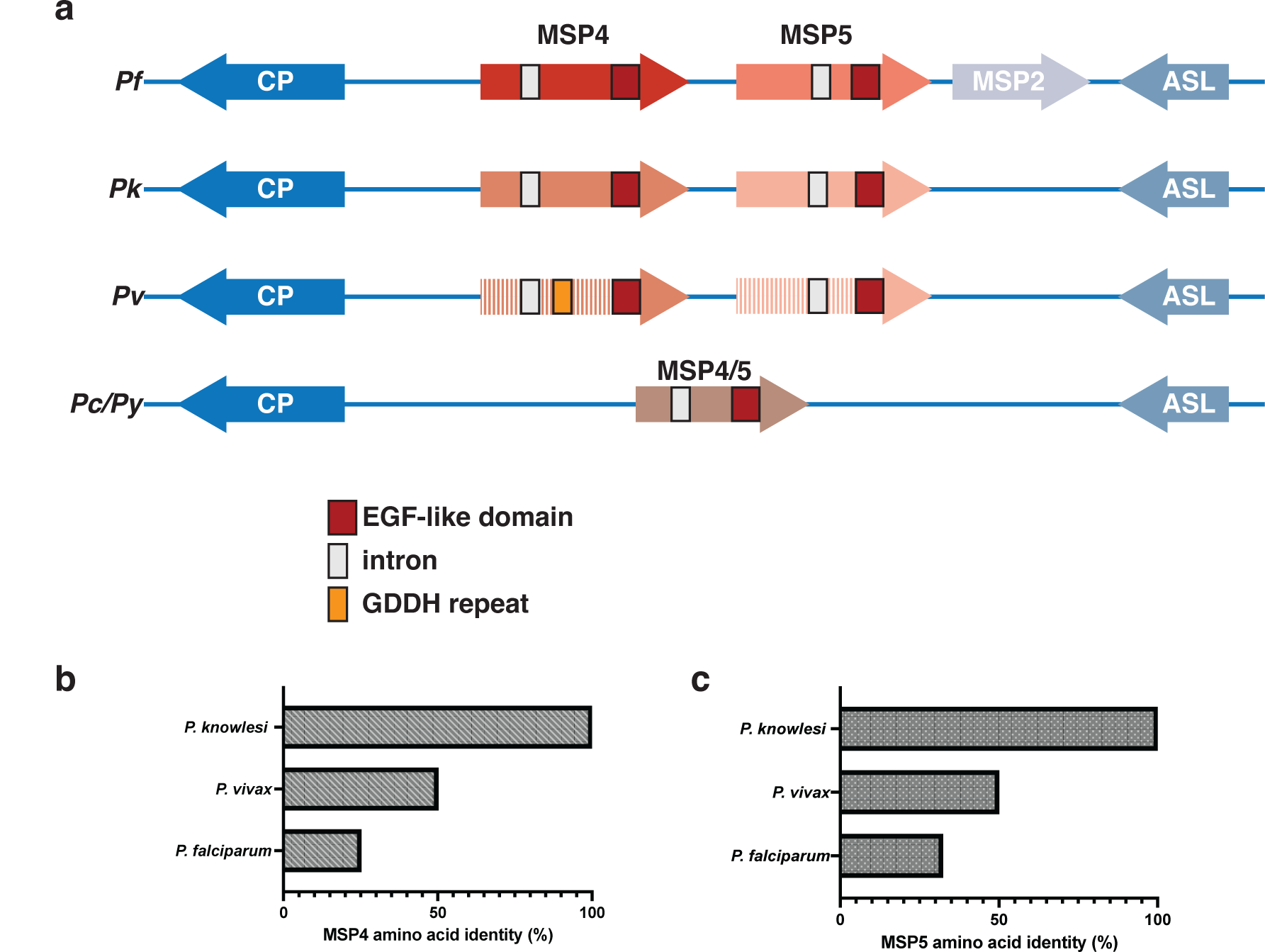
MSP4 and MSP5 exist in a syntenic region of the *Plasmodium* genome. a) Schematic representing the genetic loci and conservation of the MSP4 and MSP5 genes in human infecting *P. falciparum*, *P. knowlesi*, *P. vivax* and the rodent species *P. chabaudi* and *P. yoelii*. Also indicated are the flanking genes of this loci, an uncharacterised conserved protein (CP) and adenylosuccinate lyase (ASL). Below a graph indicating the degree of amino acid identity for b) MSP4 and c) MSP5 between the closely related species *P. knowlesi* and *P. vivax*, compared to *P. falciparum*.

Interestingly, differing levels of polymorphism are observed in the MSP4 and MSP5 genes of *P. falciparum* and *P. vivax*, suggesting they may have different functional importance. In *P. falciparum*, the average nucleotide diversity (p) for MSP4 is around ∼8.6X greater (p = 0.0031) (41) than it is for MSP5 (p = 0.00036) (42) which indicates *Pf*MSP4 is under greater selection pressure. Conversely, the average nucleotide diversity for *P. vivax* MSP5 in Colombian isolates (p = 0.03752) (43), and Iranian isolates (p = 0.06766) (44), is ∼31.3 - 56.4X greater than it is for MSP4 (isolates from various locations) (p = 0.0012) (45) indicating that for *P. vivax,* it is MSP5 which is under greater selection pressure. *P. knowlesi* MSP4 has also been shown to have little sequence diversity which suggests this also holds true for this species (46). These differences in sequence diversity indicate immune selection pressure is being exerted more so on *Pf*MSP4 and *Pv*MSP5 than *Pf*MSP5 and *Pv*MSP4.

In the present study we used Cas9 gene editing to discern the importance and function of MSP4 and MSP5 in *P. knowlesi*. We demonstrate differential importance of MSP4 and MSP5 between *P. knowlesi* and *P. falciparum* and show *P. vivax* orthologs of MSP4 and MSP5 can be expressed in *P. knowlesi*. We observe a significant invasion defect with inducible knock-down of MSP5 in *P. knowlesi*, providing direct evidence that this MSP has a role in *P. knowlesi* merozoite invasion.

## Results

### *P. knowlesi* MSP5, but not MSP4, is essential for blood stage parasite growth

Previous studies demonstrated that in *P. falciparum, pfmsp4* is essential for blood stage replication due to the inability to disrupt it using double cross-over homologous recombination (22). In the same study, *pfmsp5* was deemed non-essential as it was successfully knocked-out with no apparent change in blood stage replication. To determine whether this is also the case for *P. knowlesi*, we used spCas9 based gene editing (7, 25) to specifically disrupt *pkmsp4* and *pkmsp5.* We undertook transfection of *Pk* YH1 (6, 47) wild-type parasites with a construct to integrate an eGFP coding sequence in place of the *pkmsp4* coding sequence (Figure 2, a). As the *Pk* YH1 strain was adapted to *in vitro* culture using human RBCs by continuous culture for 12 months (6), we sequenced across the MSP4 and MSP5 coding regions which confirmed this region is identical to the annotated *P. knowlesi* strain H reference genome found on PlasmoDB. We were also able to detect MSP4 transcripts via qPCR of *Pk* YH1 derived RNA, demonstrating that *pkmsp4* is expressed in this line (Supplementary Figure 1, a). After transfection, we selected for clonal parasite populations and confirmed that wild type *pkmsp4* could no longer be detected via PCR (Figure 2, b) in 3 of 4 clones, with the fourth clone having both a wild type and integrated knock-out band which suggested a mixed culture, or by qPCR (Supplementary Figure 1, a). All parasites in the 3 confirmed transfected clones were found to express high levels of GFP via microscopy and flow cytometry (Supplementary Figure 1, b) and were positive for eGFP integration via PCR (Figure 2, b), confirming replacement of the *pkmsp4* gene with eGFP. We further characterised the 3 clonal *Pk*MSP4 KO parasites and found no change in blood stage replication when compared with the parental *Pk* YH1 line (Figure 2, c).

**Figure 2.**
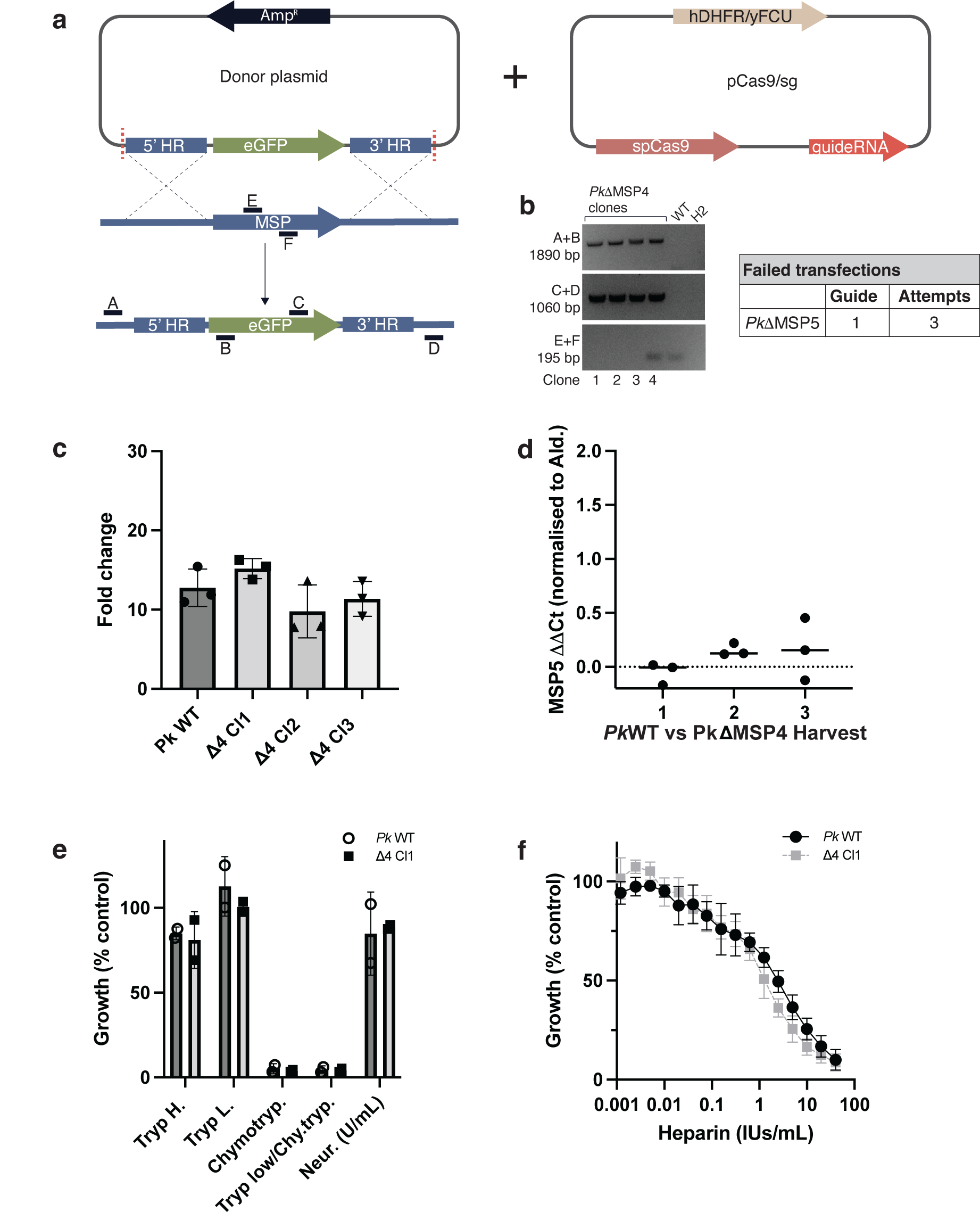
**MSP5, but not MSP4, is essential for *P. knowlesi* blood stage replication**. a) Schematic of the Cas9 gene editing strategy used for generation of *Pk*ΔMSP4 (knock-out) parasite lines. Plasmid expressing Cas9 and guide was simultaneously transfected with a linearised DNA repair template containing the eGFP coding sequence flanked by homology regions targeting *pk*msp4. b) Clonal lines were assessed via PCR for integration of eGFP using primer pairs A+B and C+D which are specific for integration and with primers E+F for wildtype (WT) pkmsp4 sequence. Multiple transfections targeting *pk*msp5 for deletion using a guide found to work efficiently for gene replacement at this loci were unsuccessful. c) MSP4 deficient parasites were assessed for changes in blood stage replication via flow cytometry and showed no change in growth compared to WT. N = 3 biological replicates, error bars are ± 1SD. d) Assessment of *Pk*MSP5 expression using qPCR. cDNA generated from 3 separate RNA harvests of *Pk*WT and *Pk*ΔMSP4 parasites was assessed via qPCR in triplicate. Ct value of MSP5 was normalised to *P. knowlesi* aldolase (ΔCt) and then WT compared to *Pk*ΔMSP4 (ΔΔCt). e) Following enzymatic treatment of RBCs, parasite replication was measured via flow cytometry following 1 replication cycle. N = 2 biological replicates, error bars are ± 1SD. f) Heparin growth inhibition assay, measured by flow cytometry following 1 replication cycle reflecting the mean ± 1SD of N = 3 biological replicates.

To determine whether *Pk*MSP5 was being upregulated in order to compensate for the lack of MSP4, *pkmsp5* transcript levels were assessed via qPCR and were found to be unchanged (Figure 2, d). There was also no change in broad receptor usage for RBC invasion as seen following enzymatic cleavage of RBC receptors (Figure 2, e) or addition of heparin (Figure 2, f). Using the same technique and a Cas9 guide targeting the *pkmsp5* locus that we demonstrated to work using a gene-replacement strategy (discussed later in the manuscript), we observed that *pkmsp5* is refractory to gene deletion (Figure 2, b). Thus, surprisingly, MSP5 appears to be essential and MSP4 non-essential for *P. knowlesi in vitro* blood stage replication. This observation is opposite to the findings of Sanders *et al*. (22) who demonstrated that MSP4, but not MSP5, is refractory to gene deletion in *P. falciparum*.

### *Pk*MSP5 and *Pk*MSP4 can be functionally replaced by their *P. vivax* orthologs

Transgenic *P. knowlesi* expressing *P. vivax* antigens have previously been shown to be useful for studying potential *P. vivax* vaccine candidates (7, 9). Therefore, we wanted to determine whether we could functionally replace *Pk*MSP4 and *Pk*MSP5 with their *P. vivax* counterparts. Cas9 gene editing was again used to simultaneously delete the *P. knowlesi* sequences and in their place integrate the *P. vivax* orthologous sequences (Figure 3, a) (Supplementary Table 1). Flanking loxP recognition sites were also included with the gene replacement to allow for future conditional knock-out (cKO) studies (48). Following transfection, we selected clonal parasite lines that were positive by PCR for *pvmsp4* integration and negative for WT *pkmsp4* (Figure 3, b top panel). When screening the *pvmsp5* clonal lines via PCR, 3 of 3 lines were positive for integration but also amplified a sequence of similar size (albeit slightly larger) to WT *pkmsp5* using primers designed to specifically amplify WT *pkmsp5* (highlighted by white box, bottom panel, Figure 3, b). These amplification products were sequenced and the results indicated it was the newly integrated *pvmsp5* sequence, and not the WT *pkmsp5* sequence, that was being amplified, confirming that these clonal lines had integrated *pvmsp5* correctly (data not shown). Importantly, the guide used to successfully integrate the *pvmsp5* into the *pkmsp5* locus was the same used in the attempts to knock-out *pkmsp5*, confirming that the guide used in the *pkmsp5* knock-out attempts was functional.

**Figure 3.**
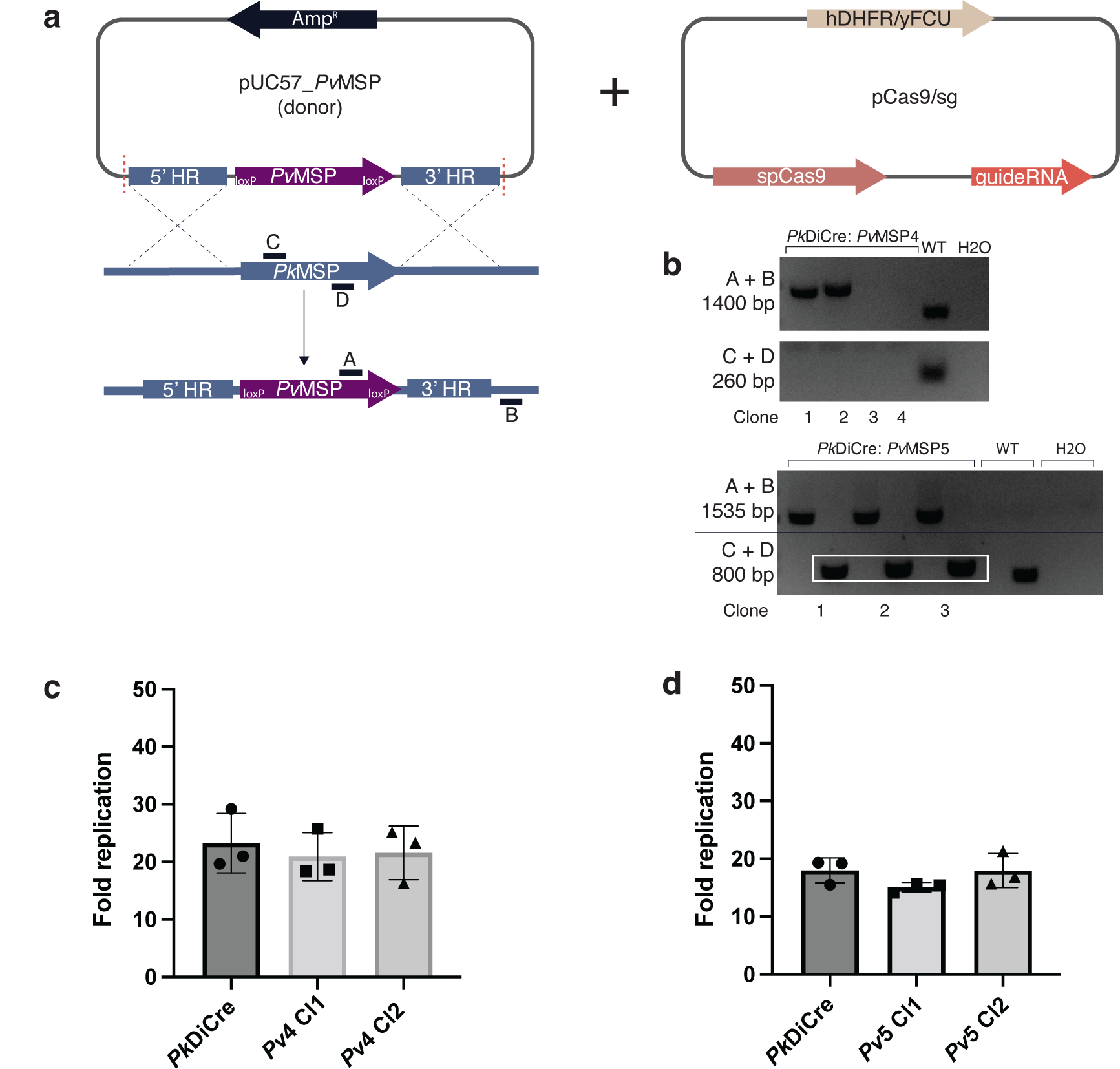
***P. knowlesi* MSP4 and MSP5 can be functionally replaced by their *P. vivax* orthologs.** a) A schematic of the Cas9 gene editing strategy used for generation of *Pk*:*Pv*MSP4 and *Pk*:*Pv*MSP5 lines. The Cas9/guide RNA expressing plasmid was simultaneously transfected with a linearised repair template which included DNA sequences homologous to the 5’ and 3’ flanking regions of the MSP being targeted and the *P. vivax* MSP coding sequence with 5’ and 3’ loxP recognition sequences. b) PCR using primers A+B allowed for detection of successful integration of the *pv*msp gene, and primers C+D were used to screen for WT *pk*msp sequence. c) *Pk*:*Pv*MSP4 and d) *Pk*:*Pv*MSP5 clonal parasite lines were assessed via flow cytometry for changes to blood stage replication. N = 3 biological replicates, error bars are ± 1SD.

The lines expressing *P. vivax* MSP4 and 5 were then assessed for any impacts on blood stage replication with the change from the *P. knowlesi* antigen to that of *P. vivax* compared to the parental *P. knowlesi* (dimerisable Cre recombinase, integrated at the p230 locus). No change in parasite growth was observed for either *Pk*:*Pv*MSP4 (Figure 3, c) or *Pk:Pv*MSP5 (Figure 3, d) when compared to the parental *P. knowlesi* line. As we had previously determined that MSP5 was essential for *P. knowlesi* blood stage replication, we conclude that the *P. vivax* MSP4 and MSP5 were being functionally expressed in *P. knowlesi*.

### Confirmation that *P. falciparum* has opposite importance for MSP4 and MSP5

Upon discovering that *P. knowlesi* MSP4 was non-essential and *Pk*MSP5 was essential, we wanted to recapitulate the findings of Sanders et al. (22) to confirm that *P. falciparum* did indeed have opposite importance for MSP4 and MSP5 compared to *P. knowlesi*. Again, Cas9 gene editing was used to target *pfmsp4* and *pfmsp5* in the parasite line NF54 and integrate in these loci an mNeonGreen coding sequence (Figure 4, a). Following transfection, we obtained clonal lines with *pfmsp5* knocked-out, as confirmed by integration of the mNeonGreen sequence at the *pfmsp5* locus (Figure 4, b) and the parasite population being positive for mNeonGreen expression via flow cytometry (data not shown). These clonal lines were also PCR negative for WT *pfmsp5* sequence, confirming *pfmsp5* had been successfully removed. Subsequent analysis of blood stage replication via flow cytometry of the *Pf*MSP5 KO clonal lines showed no change in replication when compared to the parental NF54 line (Figure 4, c) indicating *pfmsp5* is non-essential for blood stage replication. Following multiple transfections, we were unable to generate a knock-out of *pfmsp4*, confirming it is likely essential for *P. falciparum* blood stage replication. These data support the results of Sanders et al. (22) that in *P. falciparum*, MSP4, but not MSP5, is refractory to knock-out, and confirm our observation that *P. falciparum* and *P. knowlesi* show differential importance between MSP4 and MSP5 genes. Additionally, we repeated the MSP4 and MSP5 KO transfections in the Dd2 parental line, which differs from the NF54 line in its typical reliance on sialic acid residues of glycophorins on the RBC surface for invasion (49). These transfections yielded the same results as in NF54, where PCR confirmed successful deletion of MSP5 from mixed (non-clonal) transfected parasites which recovered from drug selection, but no parasites were recovered in the populations transfected with the MSP4 knock-out vectors (Figure 4, d).

**Figure 4.**
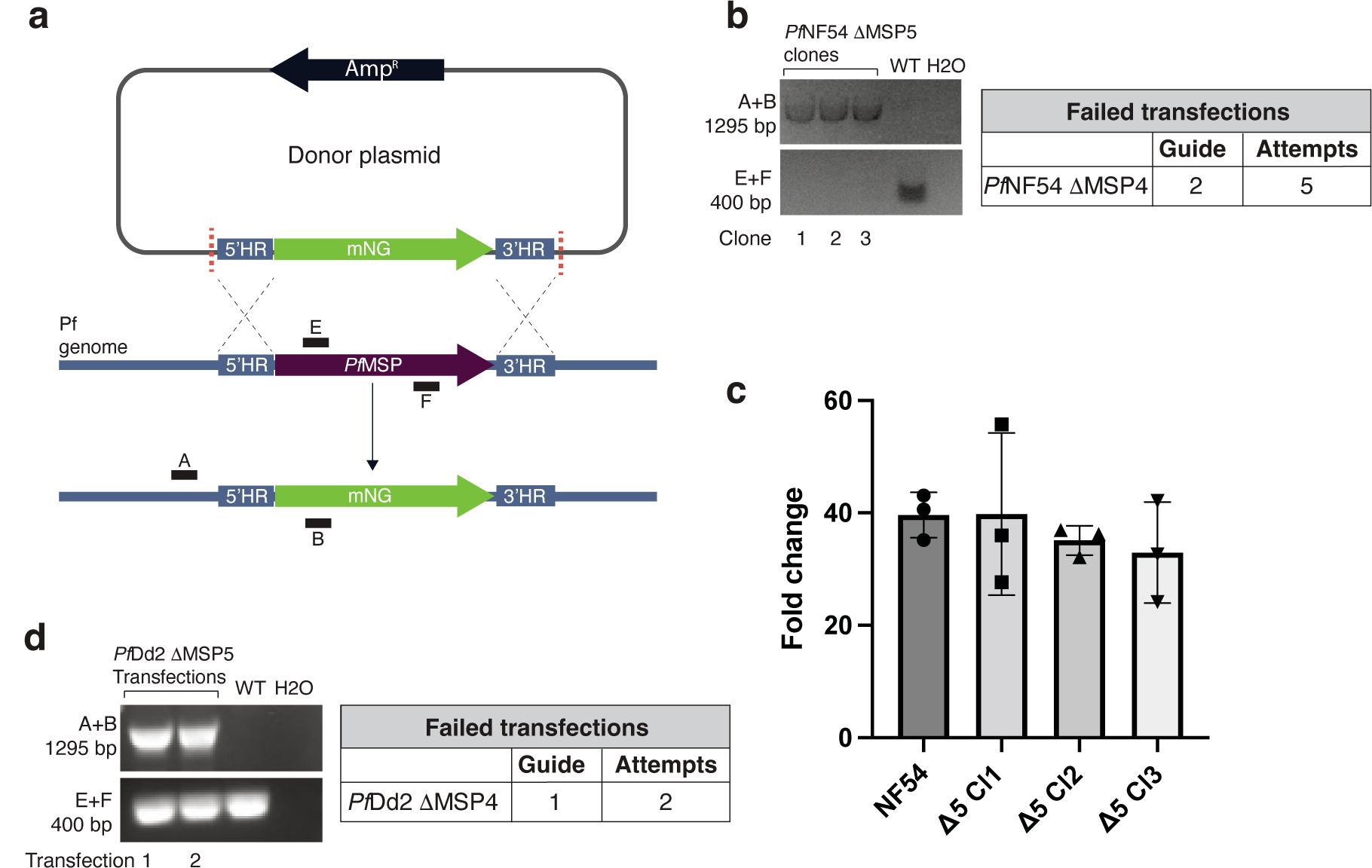
MSP4, but not MSP5, is essential for *P. falciparum* blood stage replication. a) A schematic of the Cas9 gene editing approach used to target *pf*msp4 and *pf*msp5 in *P. falciparum* laboratory isolates NF54 and Dd2. The Cas9/guide expressing plasmid was transfected simultaneously with linearised repair template that would result in replacement of the MSP sequence with the mNeonGreen (mNG) coding sequence. b) PCR using primers A+B indicated that clonal lines were positive for integration of mNG into the NF54 *pf*msp5 locus and primers E+F indicated the absence of the WT NF54 *pf*msp5 sequence. Multiple transfections aiming to knock-out NF54 *pf*msp4 using two different guides were unsuccessful. c) NF54 *Pf*ΔMSP5 clonal lines were assessed for changes in blood stage replication via flow cytometry. N = 3 biological replicates, error bars are ± 1SD. d) PCR using primers A+B indicated that two different uncloned transfected populations were positive for integration of mNG into the Dd2 *pf*msp5 locus and primers E+F indicated the presence of the WTDd2 *pf*msp5 sequence in this mixed population. Multiple transfections aiming to knock-out Dd2 *pf*msp4 were unsuccessful

### Conditional knock-out of *Pv*MSP5 and *Pf*MSP4 demonstrate their importance for blood stage replication

To further investigate the importance of MSP5 (in *P. knowlesi*) and MSP4 (in *P. falciparum*) for blood stage replication, we utilised the DiCre/loxP based cKO system (48). Here we used the *P. knowlesi* line expressing *Pv*MSP5 that can undergo cKO described above. The *Pf*MSP4 cKO line was generated in an NF54 line which expresses dimerisable Cre-recombinase (50). The guide used to insert this *pfmsp4* cKO sequence was also used in attempts to knock-out *pfmsp4*, confirming that the guide is functional and supporting that *pfmsp4* is essential for *P. falciparum* growth.

Upon addition of an analogue of rapamycin (rapalog) two subunits of Cre recombinase fuse and form the functional enzyme which then cleaves and excises the target gene between the engineered loxP sites, resulting in cKO of the loxP flanked gene. Rapalog induced excision was confirmed via PCR for both *pk:pvmsp5* (Figure 5, a, b) and *pfmsp4* (figure 5, c, d). To assess the effect of *Pv*MSP5 cKO on blood stage replication, *Pk*:*Pv*MSP5 ring stage parasites were treated with rapalog and following one replication cycle, growth was assessed via flow cytometry. The cKO of *Pv*MSP5 resulted in a ∼70% decrease in growth (Figure 5, e), confirming that MSP5 is required for *P. knowlesi* growth *in vitro*. Similarly, *Pf*MSP4 cKO ring stage parasites were treated with rapalog, and following a single replication cycle, growth was assessed via flow cytometry. We observed that cKO of *Pf*MSP4 resulted in a ∼50% reduction in growth (Figure 5, f). These data confirm the differential importance for parasite growth of MSP5 in *P. knowlesi* and MSP4 in *P. falciparum*.

**Figure 5.**
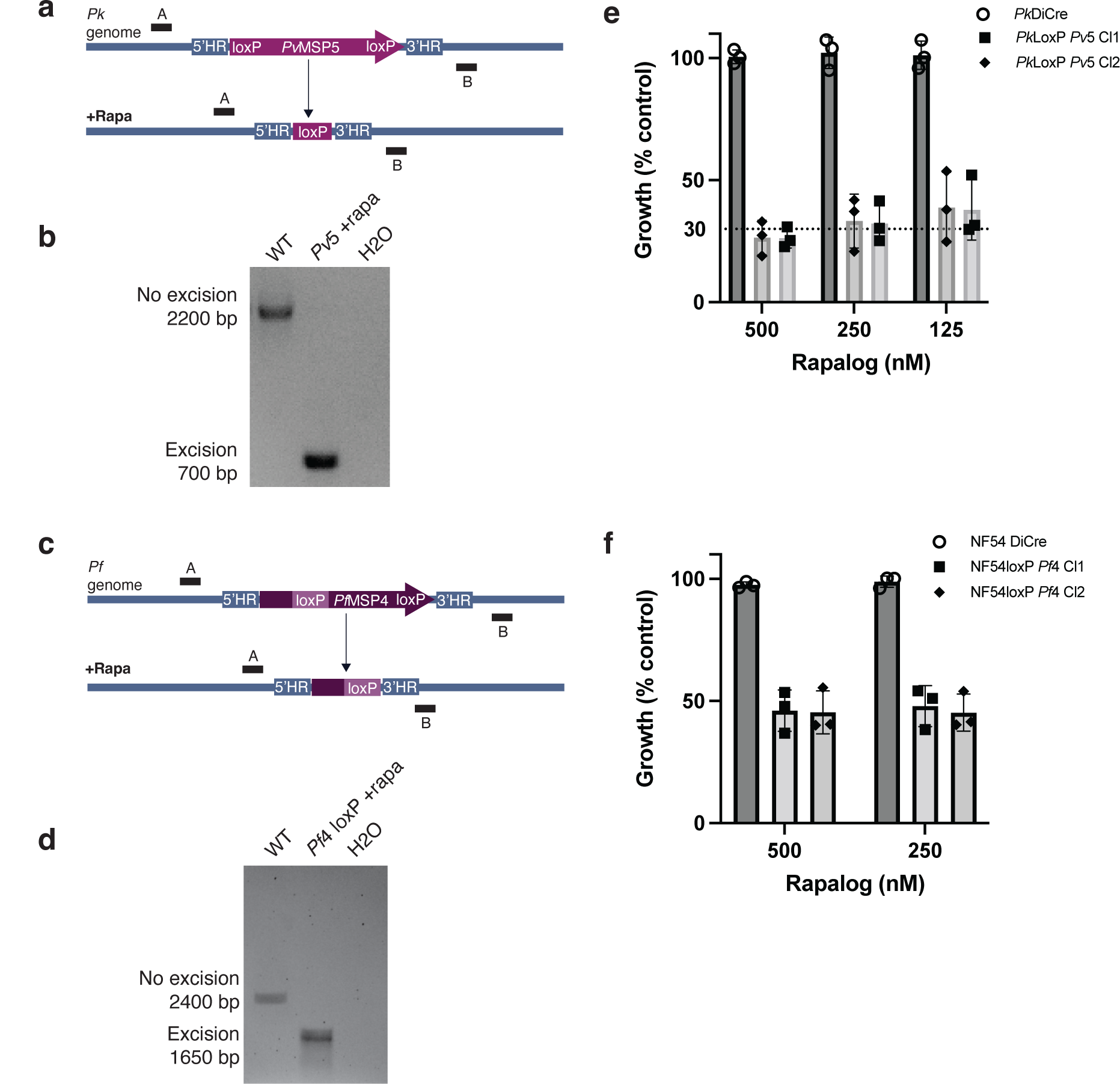
Conditional knock-out of *Pv*MSP5 and *Pf*MSP4 indicate these proteins are essential for blood stage replication. a) Schematic of the *P. knowlesi* MSP5 locus with the *Pv*MSP5 gene flanked by loxP sequences prior to and following gene excision induced by rapalog treatment. b) PCR to detect gene excision was performed using primers A+B which shows the expected ∼1500 bp decrease in fragment size following 24 hour rapalog treatment. c) The effect of *Pv*MSP5 cKO on blood stage replication was assessed via flow cytometry. *Pk*:*Pv*MSP5 parasites were treated with various rapalog concentrations and compared to untreated controls following 1 replication cycle. N = 3 biological replicates, error bars are ± 1SD. d) Schematic of the *P. falciparum* MSP4 locus with the recodonised gene, synthetic intron 5’ loxP and 3’ loxP sequences prior to and following gene excision induced by rapalog treatment. e) PCR to detect gene excision was performed using primers A+B which shows the expected ∼750 bp decrease in fragment size following 24 hour rapalog treatment. f) The effect of *Pf*MSP4 cKO on blood stage replication was assessed via flow cytometry. *Pf*MSP4 cKO parasites were treated with varying rapalog concentrations and compared to untreated controls following 1 replication cycle. N = 3 biological replicates, error bars are ± 1SD.

### *Pk*MSP5 and *Pf*MSP4 cannot be replaced by within species or across species MSP4 and MSP5 respectively

Since it was possible that conditional knock-out of *Pk/PvM*SP5 and *Pf*MSP4 could lead to selection of parasites that rely more on *Pk*MSP4 or *Pf*MSP5 respectively, we maintained *Pk/Pv*MSP5 cKO and *Pf*MSP4 cKO parasites on rapalog until cultures had returned to normal growth (Figure 6, a). Using PCR, we then tested for whether *Pk:Pv*MSP5 and *Pf*MSP4 had both been deleted from the population. However, in both cases the gene targeted for conditional deletion, *Pk:Pv*MSP5 (Figure 6, b) and *Pf*MSP4 (Figure 6, c), were still strongly fixed in the population that had grown back after extended rapalog treatment. These data support results from our gene knock-out studies that demonstrate the *Pk*MSP5 and *Pf*MSP4 are refractory to gene deletion and that *Pk*MSP4 and *Pf*MSP5 cannot readily compensate for loss of MSP5 in *P. knowlesi* and MSP4 in *P. falciparum* respectively.

**Figure 6.**
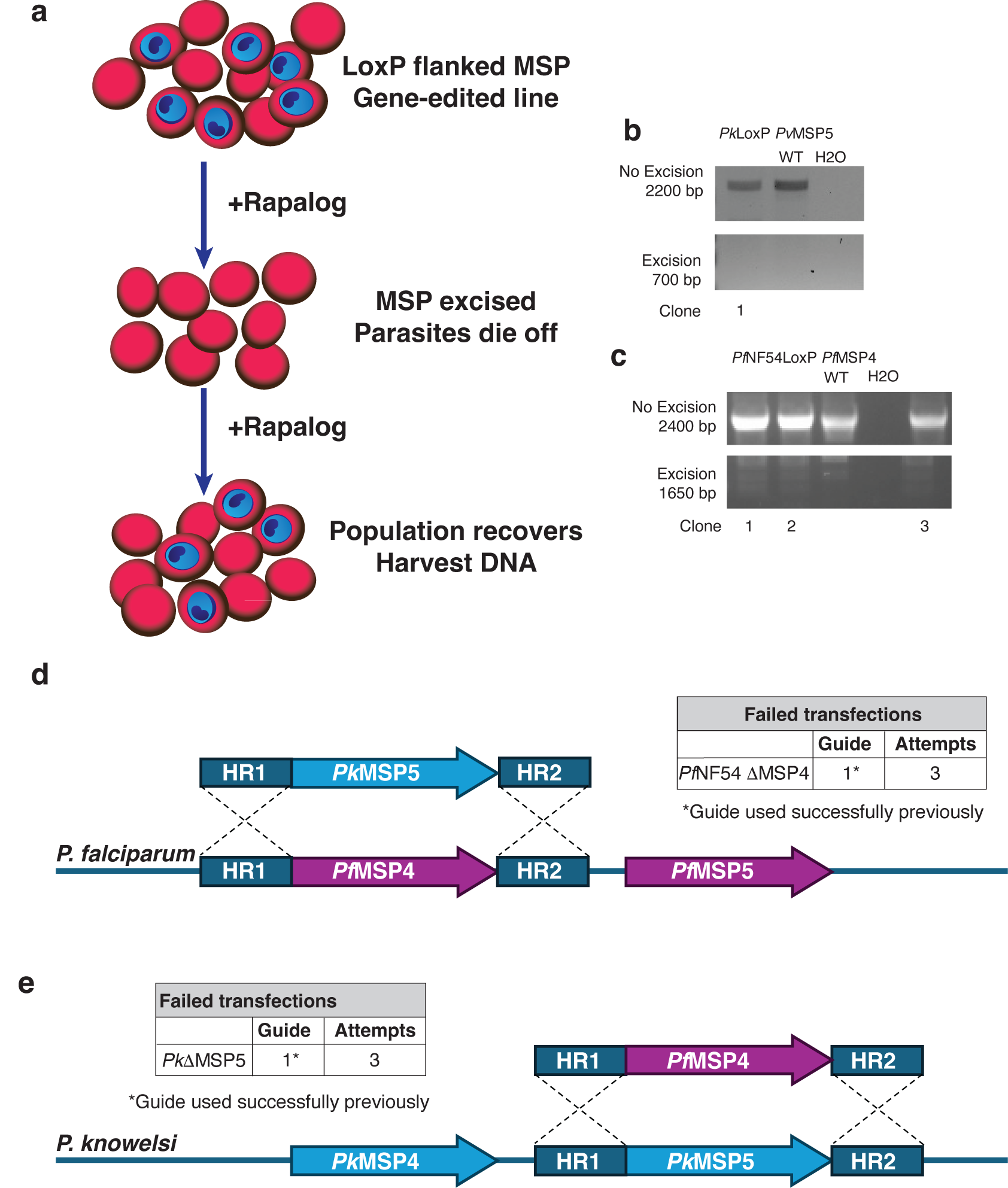
Chimeric complementation is unable to replace the function of MSP5 in *P. knowlesi* and MSP4 in *P. falciparum*. a) Schematic of the long-term target gene excision experiment where clonal LoxP flanked *Pv*MSP5 in *P. knowlesi* and *Pf*MSP4 in *P. falciparum* NF54 are excised with rapalog treatment and inducible knock-out is maintained through continuous drug pressure. Once the population recovered, parasites were harvested for DNA and the presence of the wildtype MSP or a smaller band denoting an excised gene assessed for b) *Pk*/*Pv*MSP5 and c) *Pf*MSP4. d) Schematic of the approach to replace *P. falciparum* NF54 *Pf*MSP4 with *Pk*MSP5 with the number of guides and attempts indicated. e) Schematic of the approach to replace *P. knowlesi Pk*MSP5 with *Pf*MSP4 with the number of guides and attempts indicated.

Since there was differential importance of MSP5 and MSP4 between *P. knowlesi* and *P. falciparum*, we reasoned that the essential gene from one species may be able to compensate for the loss of the essential gene from the other species. To test this possibility, we modified our CRISPR-Cas9 transfection vectors to target integration of *Pk*MSP5 into the *Pf*MSP4 locus of *P. falciparum* NF54. We also modified transfection vectors to target *Pf*MSP4 into the *Pk*MSP5 locus of *P. knowlesi*. Using the species optimised CRISPR-Cas9 gene-editing constructs and guides that we successfully used to integrate LoxP flanked *Pv*MSP5 into *P. knowlesi* and *Pf*MSP4 into *P. falciparum* NF54, we completed multiple transfections but were unable to obtain *P. falciparum* parasites that had replaced *Pf*MSP4 with *Pk*MSP5 *P. knowlesi* parasites which had *Pk*MSP5 replaced with *Pf*MSP4 (Figure 6, d) or *P. knowlesi* parasites which had *Pk*MSP5 replaced with *Pf*MSP4 (Figure 6, e). Thus, we provide evidence that the function of MSP5 in *P. knowlesi* and MSP4 in *P. falciparum* cannot be readily replaced by *Pk*MSP4 or *Pf*MSP4 and *Pf*MSP5 or *Pk*MSP5 respectively.

### Conditional knock-out of *Pv*MSP5 causes a defect in RBC invasion following initial attachment

To assess whether RBC invasion was being perturbed by cKO of *Pv*MSP5, we employed live cell microscopy to track merozoites as they attempted to invade the erythrocyte. The parental line of *P. knowlesi*, with an integrated dimerisable Cre recombinase (Pk DiCre), and the *Pk:Pv*MSP5 cKO expressing lines were kept synchronised in parallel using sorbitol lysis. Ring stage parasites were treated with 250 nM rapalog for at least 24 hours prior to filming. Invasion videos were analysed from immediately before schizont rupture until at least 5 minutes post rupture or until the final invasion event was observed. We observed the same number of merozoites develop and emerge from schizonts in the parental Pk DiCre (Supplementary Video 1) and *Pk*/*Pv*MSP5 cKO (Supplementary Video 2) lines, indicating that intracellular growth, merozoite development and rupture are generally normal despite the absence of MSP5 (Figure 7, a). cKO of *Pv*MSP5 resulted in a complete absence of merozoite invasion, whereas 35% of merozoites released from wild-type schizonts successfully completed invasion (Figure 7, b). This lost invasion potential was not due to a difference in the number of merozoites which contacted one or more RBCs (contact defined as being associated with a RBC for >2 seconds: Figure 7, c).

**Figure 7.**
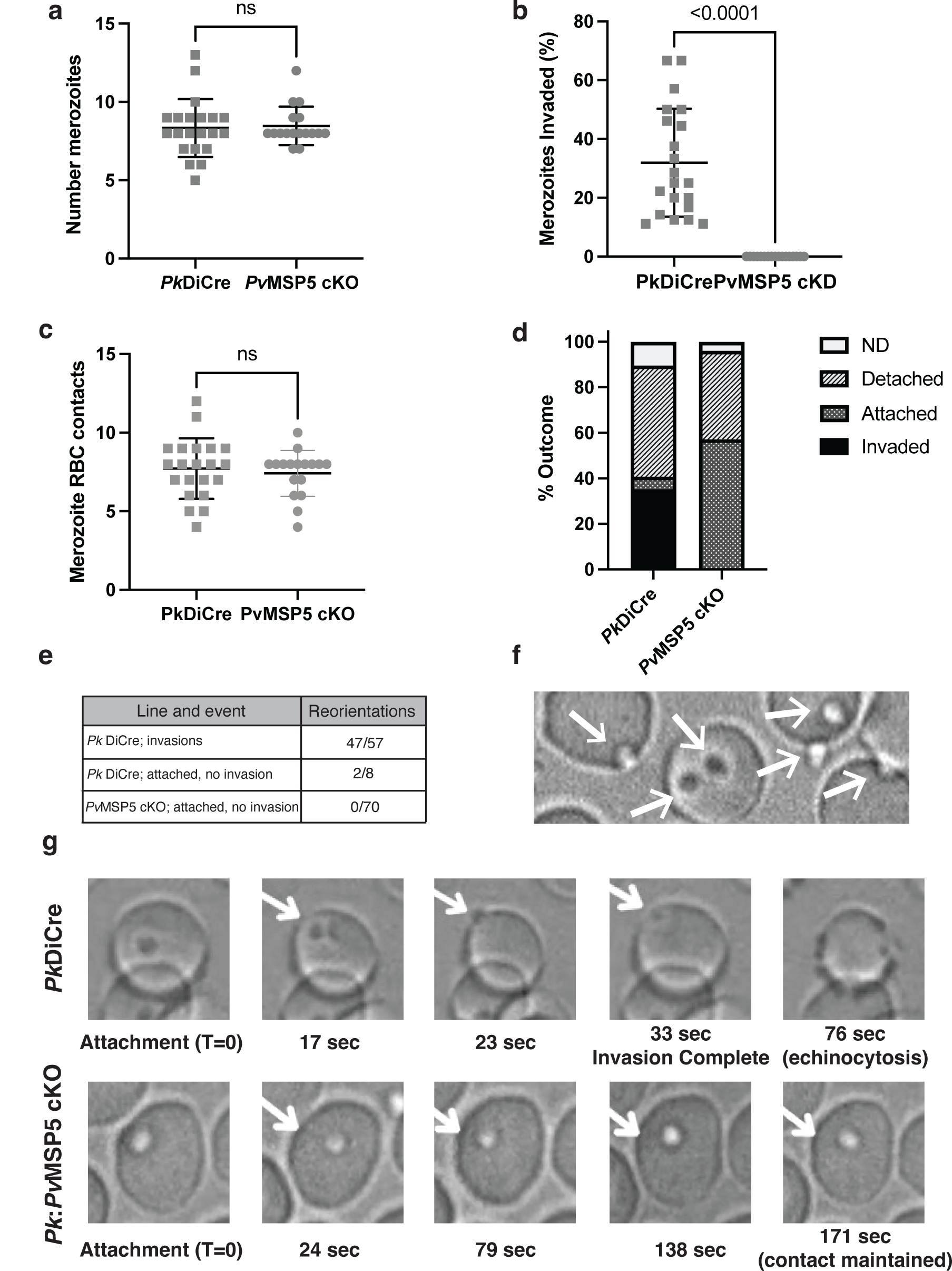
Live-cell filming reveals *P. knowlesi* merozoites deficient in MSP5 egress normally and attach to RBCs but do not invade. a) The number of merozoites each rapamycin treated *Pk*DiCre (expresses MSP5) and *Pk*DiCre *Pv*MSP5 cKO (MSP5 gene excised) schizont contained. b) The percentage of merozoites per schizont that invaded. c) The overall number of RBC contacts made by all merozoites for each schizont rupture. d) The outcomes of all schizont ruptures analysed (mean). Invaded = successful invasion event. Attached = attached and no invasion. Detached = after initially being attached, the merozoite detached and moved away from the RBC, ND = not determined (e.g. left field of view). e) The number of times clear reorientation was observable for the invasion events of the *Pk* DiCre line, for *Pk*DiCre ‘attached and not invaded’ merozoites, and for the *Pv*MSP5 cKO ‘attached and not invaded’ merozoites. f) A representative still image of *Pv*MSP5 cKO merozoites classed as ‘attached and not invaded’. g) Example still images from timepoints of a *Pk*DiCre (top panel) and a *Pk*/*Pv*MSP5 cKO (bottom panel) *P. knowlesi* merozoite after initial attachment (time post attachment under each image). Within ∼30 seconds the *Pk*DiCre merozoite had completed RBC invasion. In contrast, the *Pv*MSP5 cKO had remained attached with no progress in invasion for ∼3 minutes. For all graphs, N = 21 *Pk*: DiCre rupturing schizonts images (parental line), 17 *Pk*:*Pv*MSP5 rupturing schizonts images. All parasites were treated with 250 nM rapalog for ∼24 hours prior to filming. Error bars indicate mean ± 1SD. Statistical analysis used an unpaired T test.

Given the significant loss of invasion with *Pv*MSP5 cKO, we then analysed the steps of invasion and classed one of the following outcomes: i) ‘detach’ (merozoite moved away from, and was no longer associated with RBC), ii) ‘attached’ (merozoite remained closely associated with a particular spot on a RBC), iii) merozoite was seen to ‘invade’ and iv) ‘not determined’ (merozoite moved out of the field of view or we were unable to track it (e.g., moved under RBCs and did not reappear)). The cKO of *Pv*MSP5 resulted in an increase in the number of ‘attached’ outcomes (5.6% of merozoites in parental *P. knowelsi* merozoites compared to 57.1% for the *Pk*/*Pv*MSP5 cKO merozoites) (Figure 7, d). With *Pv*MSP5 cKO, there were no clearly observable and stable reorientation events occurring as there was with parental parasite invasion (Figure 7, e). Rather, the *Pv*MSP5 cKO parasites which were attached but could not proceed with invasion typically remained tightly associated with a single location on the RBC for an extended period of time (e.g., Figure 7, f, g). These parasites also had continued movement without noticeably reorientating or dislodging (Supplementary Video 2) indicating they had engaged with the RBC but could not proceed past this early engagement.

## Discussion

Malaria parasite merozoite surface proteins have long been studied for their potential as vaccine candidates and it has been generally assumed that these proteins function during merozoite invasion. However, recent studies have called this into question, with both *Pf*MSP1 and *Pf*MSP2 having been reported to have no role in merozoite invasion of the RBC (29, 31). Here, using CRISPR-Cas9 gene-editing and a conditional gene-deletion system we demonstrate that MSP5 is essential for RBC invasion of the zoonotic species *P. knowlesi* and that *P. vivax* MSP5 can replace *P. knowlesi* MSP5 and maintain a functional role in merozoite invasion of RBCs. We also confirmed that *P. falciparum* requires MSP4 for blood stage replication, and it is likely to play a similar role in RBC invasion. These data provide the first direct evidence that MSP4 and MSP5 have a role in merozoite invasion of the host RBC and highlight that these proteins have differential importance between malaria species.

The differential importance we found for MSP4 and MSP5 between *P. knowlesi* and *P. falciparum* is of interest given previous observations by others that the levels of polymorphism/diversity observed in clinical isolates differs between these antigens. It has been observed that *P. vivax* and *P. knowlesi* have very little MSP4 diversity when compared to *P. falciparum*, and *P. vivax* has 10-fold more genetic diversity in MSP5 than *P. falciparum* (43, 46). When designing a vaccine, it might be rational to prioritise antigens with high levels of conservation. However, in this study we demonstrate that *Pf*MSP4 or *Pk*MSP5, which are the most polymorphic of the MSP4/MSP5 antigens for each species, are also the ones where an essential function in merozoite invasion and parasite replication could be directly demonstrated. This would suggest that antibodies generated against *Pv*MSP5 and *Pf*MSP4 could be inhibiting their biological function, increasing selection pressure against them and resulting in a greater number of polymorphisms in these proteins. Whereas antibodies which target the conserved, but not obviously essential, *Pv*MSP4 and *Pf*MSP5 don’t appear to exert immune selection pressure that results in polymorphisms. These observations emphasize the necessity of understanding both the antigenicity and the biological function of potential vaccine candidates.

We demonstrate that *P. knowlesi* can functionally express *P. vivax* MSP4 and MSP5, which further promotes this system as a means for screening *P. vivax* vaccine or drug candidates. These gene edited lines provide a valuable tool to investigate the levels of protective antibodies targeting *Pv*MSP4 and *Pv*MSP5 in malaria exposed samples, as well as having potential roles in assessing subunit vaccines against these antigens. In addition, the *Pf*MSP5 KO parasites and the *Pf*MSP4 cKO parasites could similarly be used to assess protective antibody activity against these antigens in malaria exposed and vaccinated samples.

Recent *piggyBac* transposon mutagenesis studies in *P. falciparum* (51) and *P. knowlesi* (52, 53) provide a useful resource for the field when predicting likely importance of protein function in these species. The mutability index score (MIS) provided in all three studies provides the easiest way to directly compare predictions for dispensability (MIS approaching 1) or essentiality (MIS approaching 0) of genes between studies. For *Pk*MSP5, the two *P. knowlesi* saturation mutagenesis studies report conflicting findings, with one suggesting *Pk*MSP5 is likely dispensable for *P. knowlesi* A1H1(MIS 0.99) (52) and the other predicting *Pk*MSP5 is essential for *P. knowlesi* YH1 (MIS 0.16) (53). *P. knowlesi* A1H1 and YH1 are derived from the same clinical isolate, but differ in how they were adapted to *in vitro* culture in human RBCs (6, 54). Oberstaller et al. (52) identified four insertions associated with *Pk*MSP5 for A1H1, with the one mutation in the coding sequence of *Pk*MSP5 within 5.2% of the end of the gene coding region which was identified in a single sample of a possible 64 transfections. The MIS scores for *Pk*MSP4 suggest that this protein is dispensable for both *P. knowlesi* saturation mutagenesis studies (MIS 0.98 (52): MIS 0.99 (53)). However, the modified MIS+ score (0.37) preferred by Oberstaller et al. (52) led to this study assigning a tentative prediction of essentiality for *Pk*MSP4, possibly reflecting that it has only a single TTAA site towards the C-terminal end of the 791bp gene coding sequence.

Saturation mutagenesis studies in *P. falciparum* led to the prediction that both *Pf*MSP4 (MIS 0.25) and *Pf*MSP5 (MIS 0.99) are essential (51). For *Pf*MSP5 this prediction was considered tentative due to the small size of the protein and low number of TTAA sites. In contrast, both our and the study by Sanders *et al*. (22) demonstrate that *Pf*MSP5 can be knocked out of two different *P. falciparum* laboratory isolates with no impact on blood stage parasite growth *in vitro*. While our conclusions show some differences from transposon mutagenesis studies by Zhang et al. for *P. falciparum* (51) and Oberstallar et al. for *P. knowlesi* (52), this likely reflects lower confidence predictions for *Pk*MSP5 and *Pf*MSP5 due to the small size of these genes and the scarcity of TTAA sites. Overall, our findings provide direct experimental clarity on the importance of MSP4 and MSP5 to blood stage parasite growth within *P. knowlesi* and *P. falciparum*. Our study also suggests different cross-species importance between MSP4 and MSP5, supporting genetic diversity evidence where *Pf*MSP4 and *Pk*MSP5 show the most sequence diversity (42, 43, 45, 46, 55).

Although it has long been held that MSPs have a direct role in merozoite invasion, which has been supported by studies using inhibitory antibodies (16, 21, 26, 27, 56), the cKO of *Pv*MSP5 is one of the first direct demonstrations that loss of an MSP leads to significant failure to invade a host RBC. The *P. knowlesi* parasites that have *Pv*MSP5 conditionally knocked-out were able to complete intracellular growth and schizont rupture normally, and also displayed similar movement to parental *P. knowlesi* parasites in live-cell filming, indicating that conditional knock-out of MSP5 does not impact on *P. knowlesi* growth or merozoite development. Analysis of *P. knowlesi* live-cell invasion videos demonstrated that MSP5 cKO parasites remain tightly associated with a particular location on the RBC surface for upwards of 2 minutes and failed to invade the RBC, whereas the parental *P. knowlesi* parasites would proceed with invasion or dissociate from the RBC membrane well within this time. Although these parasites did not readily detach and remained tightly associated with a single location on the RBC surface, reorientation was not clearly observable, suggesting that this propensity to remain in contact with the RBC membrane is unlikely to be due to formation of the tight-junction. Instead, loss of MSP5 function in *P. knowlesi* appears to impact on parasite entry into the RBC prior to tight-junction formation.

## Conclusion

The prominent location of MSPs on the merozoite surface has supported the theory that they would have a role in RBC invasion, but there is limited direct data to support this assumption and recent studies have, instead, thrown this assumption into question. Here, we provide the clearest evidence to date that an MSP, MSP5 in *P. knowlesi*, is required for RBC invasion, with *P. knowlesi* parasites lacking this protein found to remain attached to the RBC surface for unusually extended time periods with none of the typical RBC interactions expected through this close contact. We demonstrate that MSP4 and MSP5 have differential importance between *P. knowlesi* and *P. falciparum*, highlighting assumptions about antigen importance, and potentially suitability to target with vaccines, should not be automatically ascribed from one human infecting malaria species to another. These data, and the gene-edited parasites lines produced, demonstrate that some MSPs do have important roles in merozoite invasion of the RBC and can be used to inform on assessment and prioritisation of MSP4 and MSP5 as vaccine candidates across different human malarias.

## Methods

### Parasite culture

*P. knowlesi* YH1 (6) was modified to contain dimerisable Cre-Recombinase at the pfs47 locus using CRISPR-Cas9 (7, 8) and *P. falciparum* NF54 modified to contain dimerisable Cre-Recombinase at the pfs47 locus (50). Parasites were cultured in O+ red blood cells (Lifeblood, Australia) as previously described (57). Cultures were maintained in RPMI-HEPES medium (Gibco) supplemented with either 0.5% (v/v) Albumax II (Gibco) or 0.25% heat inactivated human serum (Lifeblood, Australia)/0.25% Albumax II (Gibco), 52 mM gentamycin (Gibco), 367 mM hypoxanthine (Sigma-Aldrich), 2 mM Glutamax (Gibco) and 2 mM sodium bicarbonate (ThermoFisher Scientific) at a pH of 7.4. Cultures were incubated at 37°C within sealed boxes filled with 1% O2, 5% CO2, 94% N2.

### Plasmid generation

Plasmids constructed in this study were used as repair templates for Cas9 gene editing: *Pk*MSP4 KO, *Pk* MSP5 KO (p*Pk*CC1 backbone), pUC57 *Pv*MSP5 loxP, pUC54 *Pv*MSP4 loxP, pUC57 recod *Pf*MSP4 loxP, pUC57 *Pf*MSP4 KO, pUC57 *Pf*MSP5 KO.

These plasmids were generated by PCR amplification of regions homologous to 5’ and 3’ UTRs of the gene to be targeted (Supplementary Table 2) which were then cloned into a backbone containing: eGFP, mNeonGreen or *P. vivax* gene sequences or recodonised genes by restriction enzyme cloning. Recodonised genes and *P. vivax* sequences were synthesised by Genscript and then had homology regions cloned in or were synthesised to already contain them (Supplementary Table 1).

The Cas9, guide RNA expressing plasmids used were: pl10 (7) for *P. knowlesi* transfection and pDC2 (48) for *P. falciparum* transfection. Specific guide sequences were cloned in by first annealing single stranded oligos (Sigma-Aldrich, IDT) which contained the gRNA sequence with 15 bp of homology to plasmid for utilisation of the InFusion cloning protocol (Takara).

### Transfection

Direct electroporation of schizonts was performed as previously described (54). In brief, synchronised late stage schizonts were purified from uninfected RBCs using 70% Percoll (Cytiva) and allowed to recover at 37oC and reach full maturity by applying compound 2 (58) or ML10 (59). Repair template DNA (60 μg), linearised using appropriate restriction enzymes, and the Cas9/guide RNA expressing plasmid (20 μg) were precipitated and resuspended in TE, prior to being mixed into the ‘incomplete cytomix’ (0.895% KCl, 0.0017% CaCl2, 0.076% EGTA, 0.102% MgCl2, 0.0871% K2HPO4, 0.068% KH2PO4, 0.708% HEPES in milliQ H2O at pH 7.6). A volume of 10 - 20 µL of packed mature schizonts were resuspended in DNA/cytomix and electroporated (BioRad) before returning to prewarmed media and fresh RBCs. The day following transfection, parasite cultures were treated with either 200 mM pyrimethamine (Sigma-Aldrich) (*P. knowlesi*) or 2.5 mM of WR99210 (Jacobus Pharmaceutical Company) (*P. falciparum)* for positive selection of Cas9/guide plasmid. After 7 days, drug selection was removed and gDNA was harvested from parasite populations (Invitrogen PureLink Genomic DNA Mini Kit) and assessed via PCR for integration of desired construct. Clonal parasite lines were obtained via limiting dilution cloning (60).

### qPCR data acquisition and analysis

For analysis of mRNA expression, parasite RNA was first reverse transcribed to cDNA. The RNA sample was incubated with dNTPs (Qiagen, 40 mM) and 0.4 mg/mL random hexamer (Qiagen) and allowed to incubate at 65°C for 5 minutes and then cooled 4°C. A pre-mixed stock of the following reagents was then added to the sample: DTT (100 mM, Invitrogen), RNAsin (40 U/μL, Invitrogen), Superscript III Reverse Transcriptase enzyme (200 U/μL, Invitrogen) and 5X Superscript buffer (Invitrogen). The reaction was allowed to take place by incubating in a thermocycler with the following settings: 25°C for 5 minutes, 50°C for 60 minutes, 70°C for 15 minutes.

cDNA samples were then diluted ¼ in Ultrapure H2O for qPCR reaction. qPCR primers (Supplementary Table 3) were first confirmed to have equivalent amplification efficiencies. Reactions were set up with primers designed to target the transcript of interest and PowerUp SYBR Green Master Mix (Applied Biosystems A25742) in MicroAmp 96 well plates (using MicroAmp plate seals). The reaction was run in the QuantStudio 7 Flex System (Thermo scientific) with the following settings: 95°C for 10 minutes, (95°C for 15 seconds, 60°C for 1 minute) X40 cycles, 95°C for 2 minutes, 60°C for 2 minutes and then 2% ramp rate to 95°C for 2 minutes.

Ct values were exported and used to determine the ΔΔCt or 2-ΔΔCt values.

ΔΔCt was determined by first calculating the ΔCt of the WT and gene editing parasite lines for the gene of interest; Δ*Ct = ct (gene of interest) – Ct (housekeeping gene)* (normalising the Ct for your gene of interest in both transgenic and WT sample).

Then; ΔΔ*Ct* = Δ*CT (transgenic line) – ΔCt (wild type)*, measures the change in gene of interest expression in transgenic vs. WT sample where 0 = no change in transgenic line compared to WT. Or calculate the expression ratio; 2^!ΔΔ#$^, which provides a fold-change value where 1 = no change in transgenic line compared to WT.

### Assessment of transgenic line replication

To assess whether transgenic parasite lines had altered blood stage replication/growth, the parental and transgenic lines were first synchronised using sorbitol lysis over several replication cycles. Trophozoites – early schizonts were then adjusted to a starting parasitaemia of 0.2 - 0.4% (depending on species) at 1% Haematocrit and plated identically in two round-bottom 96 well plates in 50 μL final volume triplicates. One plate was immediately stained with 10 mg/mL ethidium bromide and analysed by flow cytometry (BD Accuri C6 plus) for starting parasitaemia, the other returned to normal incubation conditions to allow a full replication cycle to progress, prior to parasitaemia assessment by flow cytometry. Flow data was analysed using FlowJo software (version 10, FlowJo LLC). The RBC population was gated on SSC-H vs FSC-H and from this population parasitised cells were gated in PE-H vs FSC-H based on the ethidium bromide fluorescence. Fold change was determined as: fold change = (final parasitaemia/starting parasitaema).

### Conditional knock-out growth assays

To assess the effect of cKO on parasite growth, parental and transgenic parasite lines were first synchronised via sorbitol lysis and ring stage parasites were adjusted to 1% parasitaemia at 1% haematocrit in complete media. Parasite suspension (45 μL) was plated in triplicates and rapalog (Takara) was added to a final concentration of 0, 125, 250 or 500 nM in 50 μL. The plate was returned to normal incubation conditions to allow for a complete replication cycle to take place and when ∼24-hour old trophozoites were present the parasitaemia was assessed by flow cytometry and analysed as described above. The growth (% untreated control) was measured as (treated parasitaemia/untreated control parasitaemia) x 100.

### Live cell microscopy

Parasites were synchronised via sorbitol lysis and ∼3 – 5% ring stage parasites were treated with rapalog (250 nM) and kept at ≤2% Haematocrit. The following day, prior to live cell microscopy, the Haematocrit was reduced to 0.25% and volume of 200 μL was added to a chamber of a μ-Slide 8 well glass bottom coverslip (iBidi) and returned to normal incubation conditions for ∼30 minutes to allow cells to settle and reacclimatise. Microscopy was performed on a Nikon TiE using a 60X water objective and DIC. Voltage was 3 – 3.5 V and typically exposure set to 100 – 150 milliseconds. The microscope stage was enclosed within an environmental chamber heated to 37°C with humidified gas mixture set to 1% O2, 5% CO2 and 94% N2.

## Acknowledgements

The authors would like to express appreciation to Lifeblood Australia Red Cross for providing red blood donations critical to this work. This project received funding from the Australian Research Council (FT240100420 to DWW) and the Hospital Research Foundation (Fellowship (C-MCF-52-2019) and Project Grant (S-03-EOI-2021) to DWW). JC and IGH were supported by ARC-RTP PhD scholarships. We would like to thank Prof James Beeson, Dr Herbert Opi and Assoc Prof Michelle Boyle (Burnet Institute, Melbourne), Prof Robert Moon (London School of Tropical Medicine and Hygiene) and Prof Moritz Treeck (Gulbenkian Institute for Tropical Medicine, Portugal) for the insightful feedback regarding this work. Adelaide Microscopy and Dr Jane Sibbons for access and training to live-cell imaging facilities.

## Author Contributions

Conceived and designed the experiments: DWW, JC, IGH. Performed the experiments and phylogenetic analysis: JC, IGH, OR. Analysed the data: JC, DWW, IGH, OR. Paper writing: JC, DWW, IGH, OR and based on the PhD thesis of JC.

## Data Availability

Data is available from the corresponding author upon request.

---

**Supplementary Figure 1.**
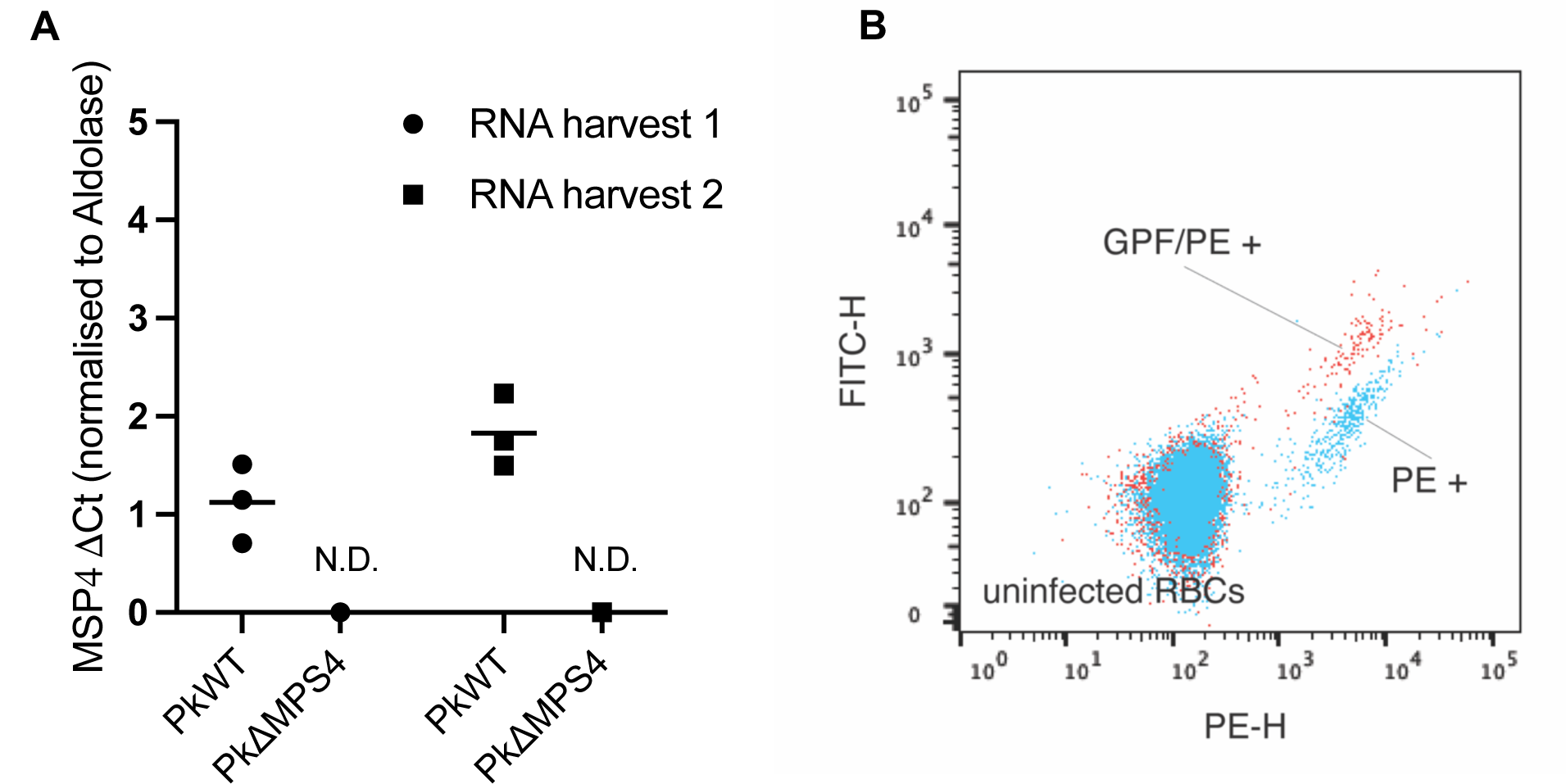
Confirmation of loss of *P. knowlesi* MSP4 expression in knock out parasites. a) Q-PCR assessment of *Pk*MSP4 expression levels in *Pk*WT and *Pk*ΔMSP4 parasites (normalised to Aldolase) across two separate RNA harvests. b) The flow cytometry data is plotted as FITC-H vs PE-H. Both the *P. knowlesi* wild-type (blue) and MSP4 KO parasite lines (red) were stained with ethidium bromide prior to flow cytometry analysis. MSP4 KO parasites are expressing eGFP as indicated by movement of population upward on FITC-H axis.

**Supplementary Video 1. Representative live cell filming video for a *P. knowlesi Pk*/*Pv*MSP5 merozoite with wildtype levels of *Pv*MSP5 invading a RBC.**

This *P. knowlesi Pk*/*Pv*MSP5 merozoite successfully invaded the RBC.

**Supplementary Video 2. Representative live cell filming video for a *P. knowlesi Pk*/*Pv*MSP5 merozoite with *Pv*MSP5 knocked out failing to invade a RBC.**

This *P. knowlesi Pk*/*Pv*MSP5 merozoite failed to invade the RBC and demonstrates the phenotype where *Pv*MSP5 inducible knock-out merozoites remain attached to the RBC for extended periods of time after attachment, but do not invade.

**Supplementary Table 1:**
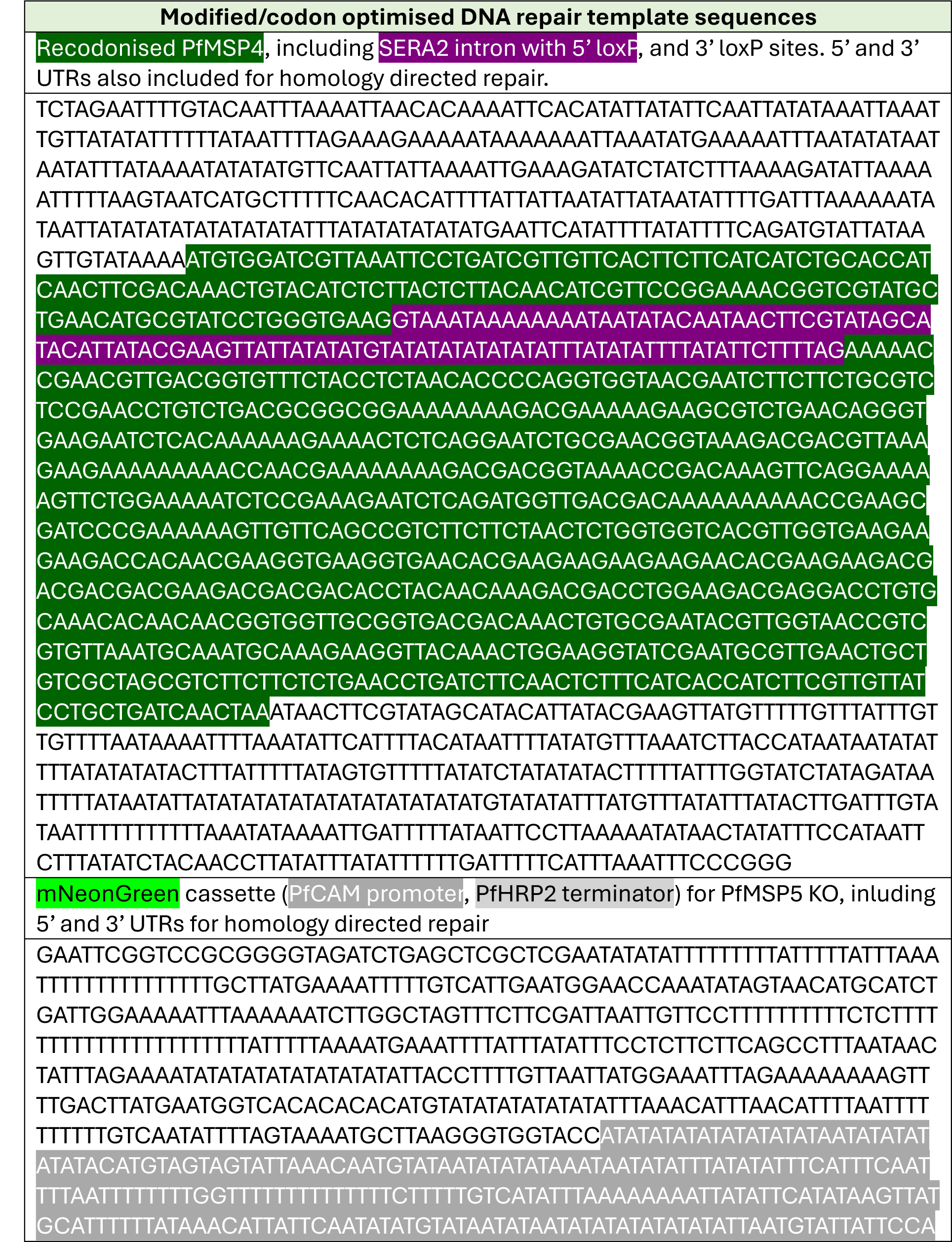
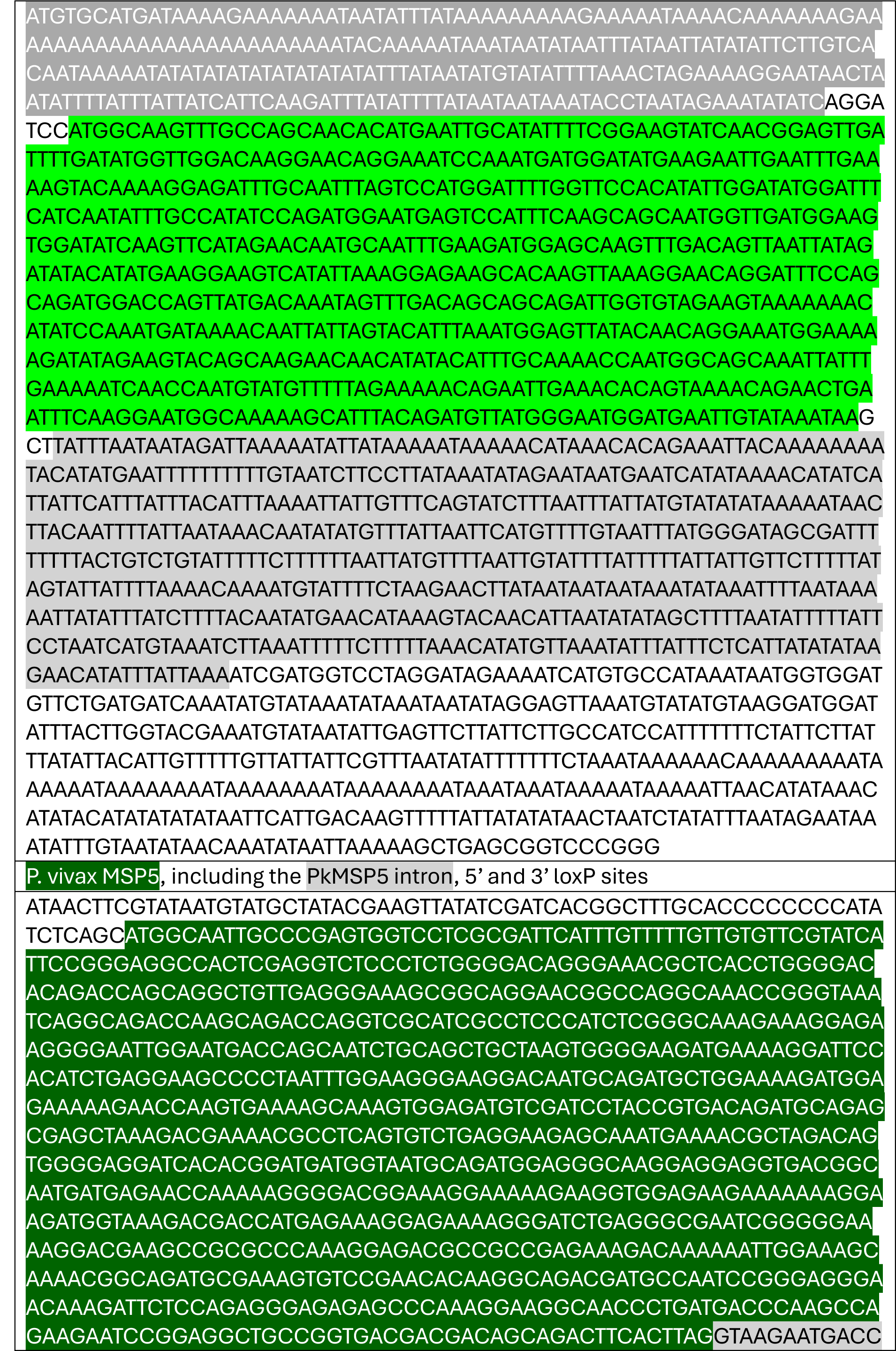
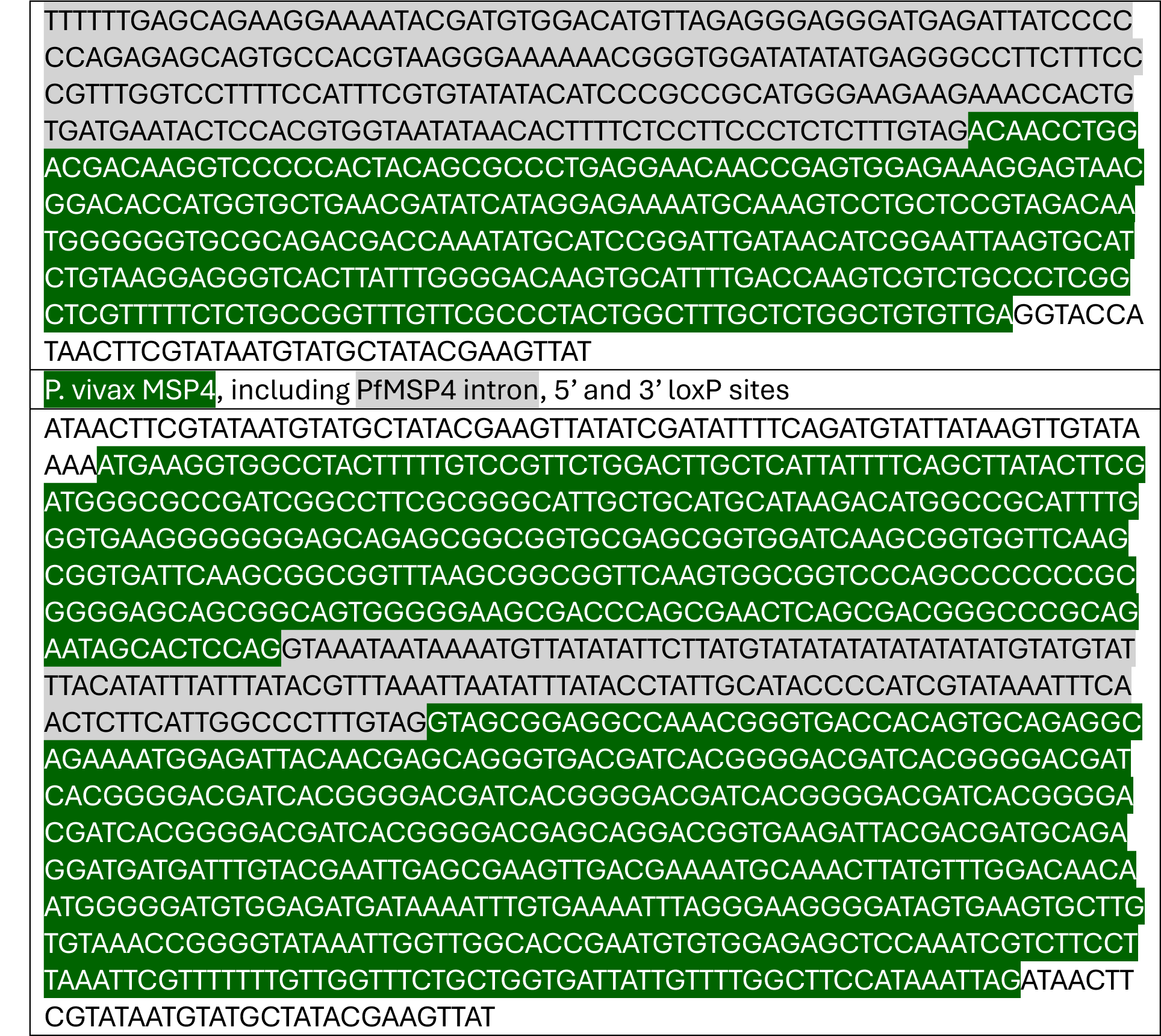
DNA sequences used for knock out or allelic replacement of *Plasmodium knowlesi* MSP4, *Pk*MSP5, *P. falciparum* MSP4 and *Pf*MSP5.

**Supplementary Table 2:**
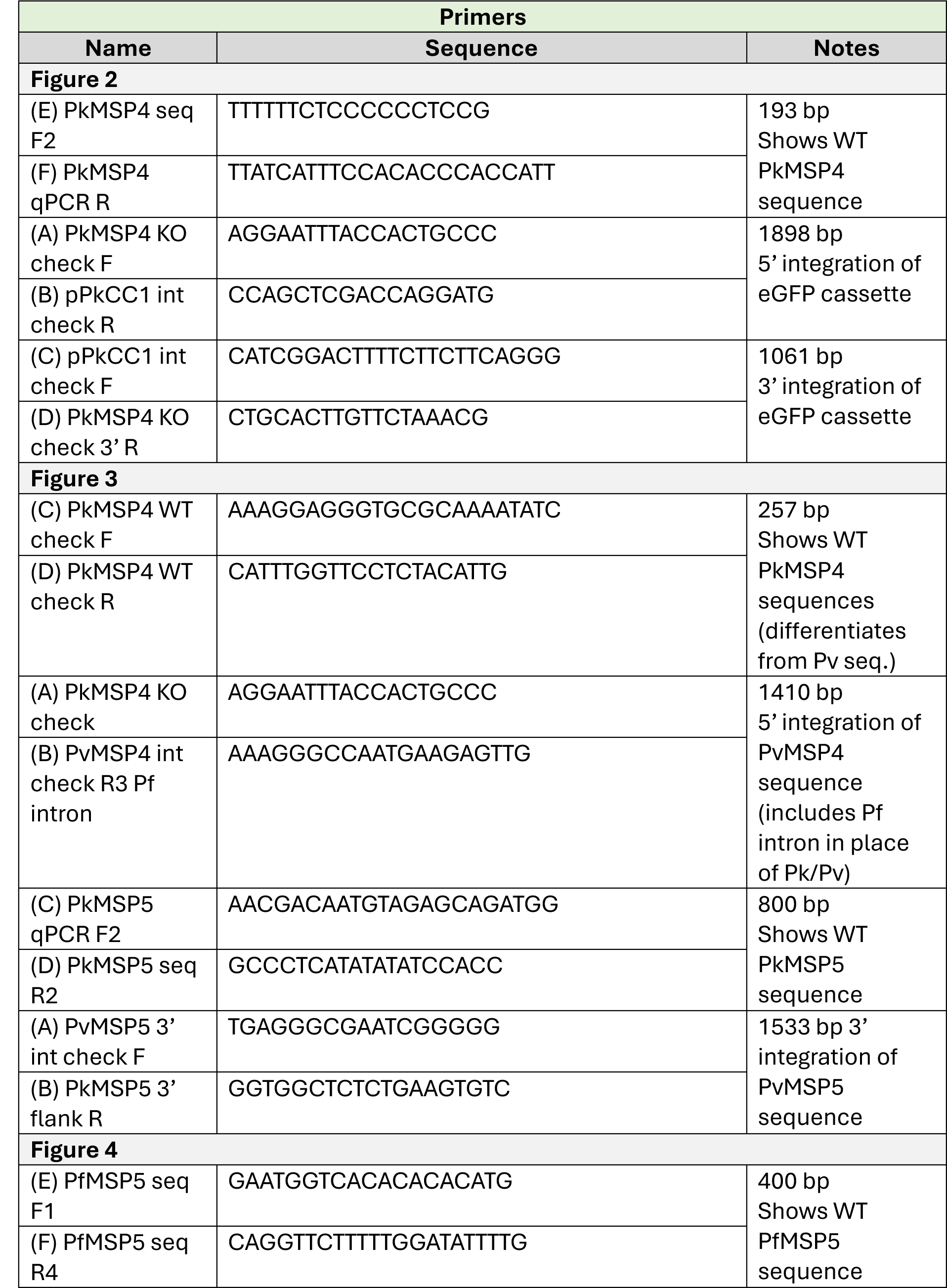
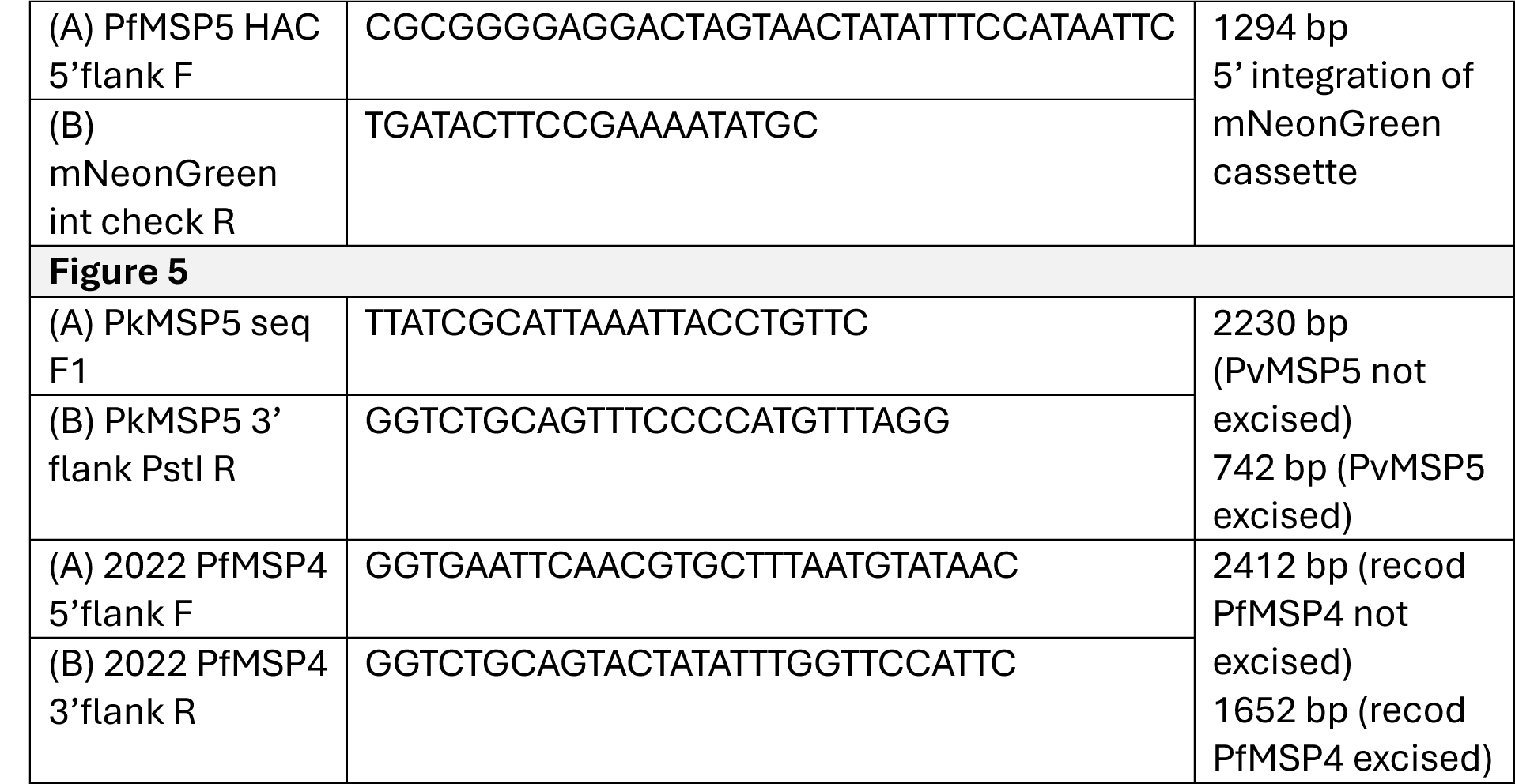
Primer sequences and expected band sizes for confirmation of *Plasmodium knowlesi* MSP4, *Pk*MSP5, *P. falciparum* MSP4 and *Pf*MSP5 gene knock outs and allele swaps.

**Supplementary Table 3:**
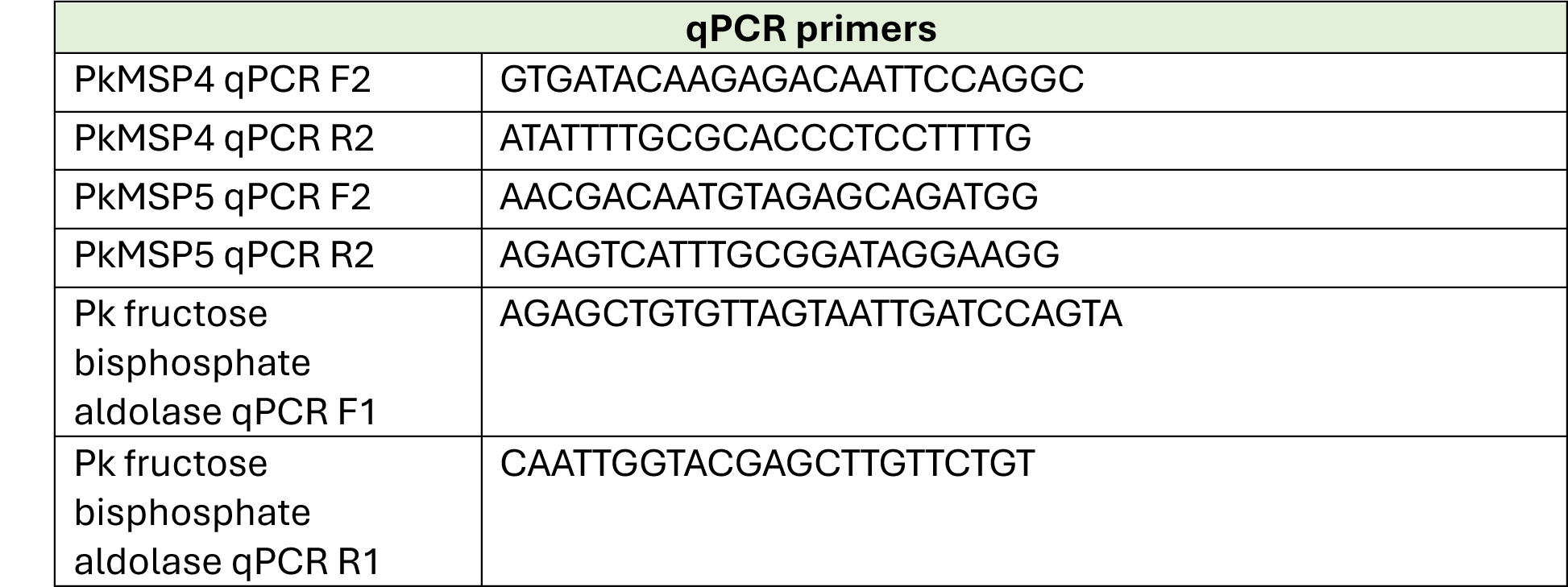
Primer sequences Q-PCR of *Plasmodium knowlesi* MSP4 and *Pk*MSP5.

